# Satellite glial cells modulate cholinergic transmission between sympathetic neurons

**DOI:** 10.1101/664557

**Authors:** Joana Enes, Surbhi Sona, Nega Gerard, Alexander C. Mitchell, Marian Haburcak, Susan J. Birren

## Abstract

Postganglionic sympathetic neurons and satellite glial cells are the two major cell types of the peripheral sympathetic ganglia. Sympathetic neurons project to and provide neural control of peripheral organs and have been implicated in human disorders ranging from cardiovascular disease to peripheral neuropathies. Here we show that satellite glia regulate postnatal development and activity of sympathetic neurons, providing evidence for local ganglionic control of sympathetic drive. We show changes in the cellular architecture of the rat sympathetic ganglia during the postnatal period, with satellite glia enwrapping sympathetic neuronal somata during a period of neuronal hypertrophy. In culture, satellite glia contribute to neuronal survival, promote synapse formation and play a modulatory role in neuron-to-neuron cholinergic neurotransmission, consistent with the close contact seen within the ganglia. Cultured satellite glia make and release neurotrophins, which can partially rescue the neurons from nerve growth factor deprivation. Electrophysiological recordings and immunocytochemical analysis on cultured sympathetic neurons show that satellite glial cells influence synapse number and total neuronal activity with little effect on neuronal intrinsic excitability. Thus, satellite glia play an early and ongoing role within the postnatal sympathetic ganglia, expanding our understanding of the contributions of local and target-derived factors in the regulation of sympathetic neuron function.

## Introduction

Glial cells, once thought of as neuron support cells, are now recognized as active players in the formation and function of normal brain circuitry [1, 2]. Astrocytes, the most abundant glial cell type in the brain, regulate many properties of neuronal circuits such as neuronal excitability, synaptic transmission and plasticity [3–5]. Their role at central nervous system (CNS) synapses has been the focus of a number of studies in the past two decades, showing that astrocytes control the formation [6–8], maturation [9], function [10, 11] and refinement [12] of synapses. These functions are mediated by various secreted as well as contact-dependent signals [11, 13, 14]. In addition to their role in the development and function of neuronal circuits [15], glia also play an important role in neurological disease, with astrocytes responding and contributing to human conditions ranging from developmental to degenerative disorders and traumatic lesions [16, 17].

In contrast to the wealth of information available on the roles of CNS astroglia, we have only a limited understanding of the satellite glia found in peripheral ganglia. This is particularly true for the sympathetic nervous system, which innervates most internal organs and regulates their function. A basal level of sympathetic activity, or sympathetic tone, together with opposing activity from the parasympathetic nervous system, ensures constant bodily homeostasis. Sympathetic tone may rise on a short timescale in response to a physiological demand (for example, exercise or stress) [18, 19], or over a long timescale, in a sustained manner, under pathological conditions such as hypertension and chronic heart disease [20, 21]. Sympathetic tone is initially set by neurons present in the brain and spinal cord [22], with the sympathetic ganglionic neurons acting as the final regulatory element determining the output of the sympathetic circuit.

A striking anatomical feature of the sympathetic ganglion is the presence of satellite glia that form an envelope around individual ganglionic neuronal somata and cover synapses [23]. This is in contrast to the CNS where individual astrocytes are in contact with multiple neurons [24]. While the function of the satellite glia remains to be fully defined, both sympathetic and sensory satellite glia share several cellular and molecular features with astrocytes, including expression of neurotransmitter receptors and the formation of a glia network via gap junctions [25]. Satellite glia injury responses are characterized by changes in expression profiles, including an up-regulation of the activation marker glial fibrillary acidic protein (GFAP) [26]. These findings point to a possible effect in disease progression and suggest that satellite glia play roles in both normal function and disease in the peripheral nervous system.

Recent studies using genetic manipulations of sympathetic satellite glia have implicated these cells in the regulation of target organ function by demonstrating that selective activation of Gq-GPCR (G protein-coupled receptor) signaling in peripheral glia leads to the modulation of cardiac properties in adult mice [27, 28]. These effects are mediated through postganglionic sympathetic innervation of the heart raising the possibility that activated glia influence the active properties of sympathetic neurons within the ganglia. This idea is supported by the finding that ganglionic cells can alter the short-term plasticity of single sympathetic neurons cultured in isolated conditions [29]. Less is known however, of the effects of satellite glia on the formation and function of cholinergic synapses in the sympathetic system.

Developmentally, reciprocal interactions between embryonic sympathetic neurons and presumptive glial progenitors in the local ganglionic environment have been shown to promote co-differentiation of both cell types at early development times [30]. Work showing that non-neuronal ganglionic cells support the early development of dendrites [31] and transiently regulate the expression of potassium currents during the perinatal period [32, 33] suggests that satellite glial cells in the sympathetic ganglia might also regulate the emergence of mature neuronal properties of sympathetic neurons. Thus, neuron-glial interactions in the sympathetic ganglia may be established early and regulate multiple properties of sympathetic neurons during the establishment of the sympathetic circuits.

Here we show that satellite glia surround sympathetic neurons within the postnatal sympathetic ganglion during a period of neuronal maturation. We explore the influence of these glia on sympathetic neuron survival and synaptic development during the neonatal period. We find that satellite glia promote the survival of cultured sympathetic neuron via a nerve growth factor (NGF)-dependent mechanism, suggesting glia contributions to neuron survival that are independent of known target-dependent survival pathways. Co-cultured glia also increase sympathetic cholinergic synaptic activity via a mechanism that involves secreted factors and regulation of the number of synaptic sites. These experiments provide insight into the development and modulation of sympathetic tone at the sympathetic-cardiac circuit’s last neuron-to-neuron synapse (see Fig 1).

**Fig 1.**
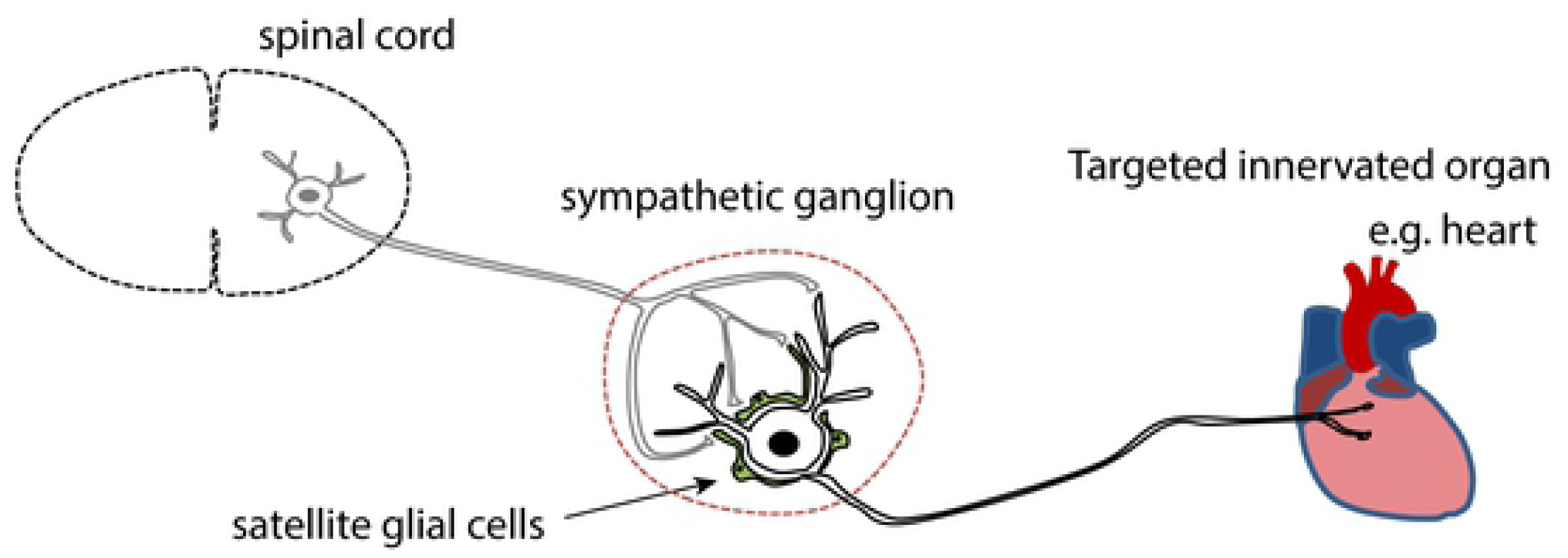
Schematic of the peripheral sympathetic-cardiac circuit. Within the sympathetic ganglia, pre-synaptic inputs from spinal cord preganglionic neurons form cholinergic synapses onto postganglionic sympathetic neurons, and satellite glial cells in the ganglia enwrap neuronal soma. The postganglionic neurons project to peripheral targets including the heart.

## Materials and methods

### Cell culture

All experimental procedures involving animals were approved by the Brandeis Institutional Animal Care and Use Committee. Superior cervical sympathetic ganglia (SCG) were dissected from P1-P3 Sprague-Dawley rats unless otherwise stated, de-sheathed, and incubated at 37° C for 1 hour in minimum essential medium (Gibco BRL, Invitrogen, Carlsbad, CA, USA) containing 350 units/ml collagenase type I (Worthington Biochemical Corporation, Lakewood, NJ, USA) and 5.5 units/ml dispase (Gibco BRL, Invitrogen, Carlsbad, CA, USA). Following enzymatic digestion, the cells were dissociated by passing repeatedly through fire-polished glass pipettes, and pre-plated on uncoated plastic tissue culture dishes for 1 hour at 37° C to remove non-neuronal flat cells. The less adherent cells, which consisted of aggregates of neurons and satellite glia, were then rinsed off the dishes, and plated at a density of 10,000 cells per dish on glass-bottomed plates (MatTek Corporation, Ashland, MA, USA) coated with collagen (50 µg/ml; BD Biosciences, Bedford, MA, USA) and mouse laminin (5 µg /ml; BD Biosciences, Bedford, MA). Cultures were maintained in modified L15CO_2_ medium (Hawrot and Patterson, 1979; Lockhart et al., 1997), supplemented with 10% fetal bovine serum (Omega Scientific, Tarzana, CA, USA), 6 μg/ml dextrose, 2 mM glutamine (Invitrogen, Carlsbad, CA, USA), 100 U/ml penicillin & 100 μg/ml streptomycin (Invitrogen, Carlsbad, CA, USA), 1 μg/ml 6,7, dimethyl-5,6,7,8-tetrahydropterine (DMPH4, Calbiochem, San Diego, CA, USA), 5 μg/ml glutathione (Sigma, St. Louis, MO, USA) and 100 μg/ml L-ascorbic acid. Mouse 2.5S NGF (5 ng/ml, BD Biosciences) was added to all cultures, unless stated, to support neuronal survival. Half of the media was exchanged with fresh NGF-containing growth medium three times weekly. To obtain glia-free neuronal cultures, cytosine arabinofuranoside (AraC, 1 μM, Sigma, St. Louis, MO, USA) was added to the cell culture media from day 1 to day 3 to inhibit glia cell division. To obtain neuron-glia co-cultures, AraC was withheld from the media and satellite glial cells proliferated rapidly, reaching 100% confluency at around 7-10 div (days in vitro). In some experiments we first used AraC to obtain glia-free neuron-alone cultures and satellite glial cells were re-plated on top of the neurons after 7 day of culture at approximately 100,000 glial cells per dish (NG[7]). These glia also formed a confluent layer by 10-14 div. Under all of these culture conditions, about 95% of the non-neuronal cells stained positive for S100β, a glial-cell marker. For NGF deprivation experiments, cultures were initially plated in the presence of 5 ng/ml NGF in serum-free medium, which was replaced after two days with NGF-free, serum-free medium. NGF-free cultures were treated with anti-NGF antibody (1:1000, final concentration 1 µg/ml), or K252 (1:20000, final concentration 100 nM).

### Immunohistochemistry

Wistar Kyoto (WKY) rats were euthanized by CO_2_ asphyxiation and SCG were dissected. The tissues were fixed for at least overnight in 4% paraformaldehyde (PFA) and then cryo-protected by incubating them in 30% sucrose solution at 4°C until the tissues sank. The tissues were placed in cryo-molds and embedded in O.C.T. (optimal cutting temperature) compound (Tissue-Tek O.C.T. Compound, Sakura Finetek, VWR, CA, USA) before freezing with dry ice.

The tissues were cut into 10 μm, longitudinal sections in a cryostat (Leica CM3050, Buffalo Grove, IL, USA) and thaw mounted onto Fisherbrand™ ColorFrost™ Plus Microscope Slides. The tissue sections were rehydrated in PBS before treatment with 10 mg/ml sodium borohydride solution and then incubated in 3% bovine serum albumin (BSA)/0.3% Triton X-100 solution for 1 hour. They were then incubated overnight with primary antibodies at the following concentrations: chicken anti-Microtubule Associated Protein 2 (MAP2) polyclonal antibody (Sigma-Aldrich, EMD Millipore, Darmstadt, Germany, AB5543, 1:1000) and rabbit anti-S100 calcium-binding protein B β-subunit (S100-β) polyclonal antibody (Agilent Dako, Santa Clara, CA, USA Z0311, 1:400). Following washing, they were incubated with donkey anti-chicken rhodamine and donkey anti-rabbit Alexa 488 secondary antibodies for 1.5 hours and then with 1 mg/ml 4’,6-diamidino-2-phenylindole (DAPI, Invitrogen Life Technologies) (1:20) for 15 mins. Subsequently, the slides were immersed briefly in distilled H_2_O and then mounted using 1:1 glycerol:PBS mounting solution. The sections were then imaged using the Zen software (Zeiss) on a Zeiss LSM 880 laser scanning confocal microscope.

### Cell density and morphology quantification

Three SCG sections per animal and 2-4 images per section were taken using the 561 nm, 488 nm and 405 nm lasers to excite the three fluorochromes: rhodamine, Alexa 488, and DAPI, respectively. Neurons in SCG sections were identified by MAP2 staining, glial cells by S100β staining, and nuclei by DAPI staining. The number of neurons was counted using the Cell Counter plug-in of the Fiji (SciJava Consortium) software. Cells stained for both S100β and DAPI were identified as glial cells and the number of glial cells was calculated. The neuron soma size was measured by manually outlining MAP-2 stained neurons within a rectangular area of identical size and position in each image using Fiji software in sections stained for MAP2, S100β and DAPI.

### Electrophysiology

Neuronal whole-cell patch-clamp recordings were made using an Axopatch 200B amplifier (Axon Instruments, Union City, CA, USA). Extracellular solution contained, in mM: NaCl 150, KCl 3, MgCl_2_ 2, HEPES 10, CaCl_2_ 2 and D-glucose 11; pH 7.4 and adjusted to 320 mOsm with sucrose. Patch pipettes had resistances of 2-4 MΩ and were filled with internal solution containing, in mM: K gluconate 100, KCl 30, MgSO_4_ 1, EGTA 0.5, HEPES 10, K_2_ATP 2, NaGTP 0.3, Tris phosphocreatine 10; pH 7.2 and adjusted to 290 mOsmol with sucrose. All recordings were made at 33-35° C using a QE-1 heated culture dish platform (Warner Instruments Inc., Hamden, CT, USA). Data were acquired with pClamp 8 software suite and digitized at 10 kHz and low-pass filtered at 2 kHz. Electrophysiological responses were analyzed using built-in functions in MatLab (The MathWorks, Inc.).

Spontaneous activity was recorded for 5 minutes at a holding potential of −60 mV; cells were classified as silent if they showed fewer than 10 single events in the 5 min period. Total synaptic charge was defined as the area above the curve, i.e. the sum of all the current values above a threshold of 25 pA. Average synaptic charge corresponds to values calculated per 10 s duration. Values presented in plots are average membrane currents quantified as averaged synaptic charge normalized to 1 ms duration. Due to incomplete voltage clamp, we occasionally found cells that showed escaping action potentials identifiable based on an amplitude > 1 nA and duration < 7.5 ms. Those spikes were excluded from the quantification by cutting them off from the original trace and replacing them by interpolated values. Series resistance (R_s_) was monitored throughout recordings but not compensated. Cells were accepted for analysis only if they met the following criteria: a) resting Vm < −45 mV, b) R_series_ < 20 MΩ, c) R_input_ > 100 MΩ and not varying more than 20% of the initial value over the course of the recording.

Evoked activity was recorded in normal extracellular solution containing the nicotinic cholinergic antagonist hexamethonium bromide (100 μM; Sigma, St. Louis, MO, USA). A small dc current was injected to maintain membrane potential at −60 mV in between depolarizations. To examine the firing properties, incremental current pulses of 500 ms duration were injected into the cell. The average cell response was calculated from 3 consecutive trials.

### Immunocytochemistry

Cultured cells were fixed with 4% paraformaldehyde and stained for ß-tubulin class III with ms anti-Tuj-1 antibody (Covance; 1:2000), for the glial cell marker S100β with rb anti-S100 (Agilent Dako; 1:1000) and for nuclei with DAPI (4’,6-Diamidino-2-Phenylindole Dihydrochloride; Invitrogen; 1:500). The antigen-antibody complex was visualized using the secondary antibodies dk anti-ms Rhodamine (1:500) and dk anti-rb FITC (1:500). Synaptic puncta were identified by the co-localization of pre-synaptic Vesicular Acetylcholine Transporter (VAChT) protein and the post-synaptic Shank protein in MAP2 stained neurons using rb anti-VAChT (Sigma Aldrich; 1:1000), ms anti-shank (Neuromab; 1:200) and ck anti-MAP2 (Chemicon, 1:1000) primary antibodies in conjugation with 1:500 diluted gt anti-rb Alexa 647, gt anti-ms Alexa 488 and gt anti-ck Alexa 568 secondary antibodies (Invitrogen).

### Synapse quantification

Sixteen bit images of 15-30 isolated neurons across 2 coverslips per condition from 3 independent cultures were acquired using a Leica DM6000 Confocal microscope under a 63x oil objective at zoom 3 and 1024×1024 resolution. Images were acquired sequentially under identical settings of laser strength, detector gain and detector offset across all conditions within each culture. These settings were chosen to exclude signal saturation in each channel using Quick Lookup Tables (QLUT) available in the Leica image acquisition software. The maximum intensity projection of each image was then analyzed using Puncta Analyzer (an ImageJ plugin written by Barry Wark and available upon request from). The number and size of synaptic puncta on SCG neuronal cell bodies and proximal dendrites (<50 μm) were quantified using identical threshold values for all cells in both conditions. The number of synaptic puncta was normalized to the MAP2-positive area. Size of synaptic puncta was calculated by the ImageJ plugin.

### Preparation of glial cell-conditioned medium (GCM) and control medium (C)

Ganglion cells were grown in serum- and NGF- containing media until confluence, about 7-9 days. The cells were then trypsinized and transferred to new 10 cm dishes. After 20 minutes, cells were washed 3 times with warm PBS to remove neurons, and cultured in serum-free NGF-free medium for 3 additional days. This glial cell-conditioned medium (GCM) was collected, centrifuged for 3 min to pellet cell debris, and concentrated using centrifugal concentrators (Sartorius) with a size cut-off filter of 5 kDa. By centrifuging at 1750xg for 90 minutes, GCM was concentrated to about 20x. GCM was then filtered through a 0.22 μm syringe filter and stored at −20°C. It was added to the cells at 1:3 ratio in fresh serum-containing media (final serum concentration of 7.5%). Control, unconditioned media (C) was prepared by concentrating approximately 20x serum-free media using the same centrifugal concentrators, and added to the cells at a 1:3 ratio in fresh serum-containing media (final serum concentration is also 7.5%).

### Western blotting

Cultured glial cells were lysed in RIPA buffer, and protein concentration in the lysate was determined using the Bradford Assay. SDS-PAGE was performed using 20 μg of each sample. The proteins were then transferred to polyvinylidene fluoride (PVDF) membrane (0.2 μm, Bio-Rad). The membrane was blocked for 1 h at room temperature using 10% non-fat dry milk in PBS, incubated with the appropriate primary antibody for 2 hours at room temperature: rb-NGF (1:500; Santa Cruz #sc-548), rb-BDNF (1:500; Santa Cruz #sc-546) or rb-actin (1:7500; Odyssey # 92642210), washed and then incubated with the respective secondary antibody (gt anti-m HRP (1:7500; Jackson ImmunoResearch #111035144) or gt anti-rb HRP (1:7500; Jackson ImmunoResearch #111035144) for 1 hour at room temperature. Both primary and secondary antibodies were diluted in 1% PBST. Blots were developed using LumiGLO Chemiluminescent Substrate (KPL# 546100) on Blue Devil X-ray Films (Genesee Scientific #30-100). To test for specificity of the antibodies, 95% confluent HEK cells were transfected either with empty (-), NGF-expressing (NGF) or BDNF-expressing (BDNF) plasmids. The NGF and BDNF constructs were cloned into promoter based SRα-based expression vector pBJ-5 [34, 35] and kindly provided by Masami Kojima (National Institute of Advanced Industrial Science and Technology, AIST Kansai, Osaka, Japan).

### Statistics

Data from at least 3 independent sets of cell culture experiments and at least three animals for immunohistochemistry were pooled for analysis. Results are presented as mean ± s.e.m.; for cell culture experiments n represents the number of neurons analyzed; for immunohistochemistry n represents number of animals. Statistical analysis was done using SigmaStat or IBM SPSS software. t-tests or Mann-Whitney were used for comparisons. For multiple comparisons, ANOVA was used, followed by pairwise post hoc (Tukey’s HSD) comparisons.

## Results

### Dynamic changes in the ganglionic structure of sympathetic neurons and satellite glia during the postnatal period *in vivo*

Satellite glia within the peripheral sympathetic ganglia enwrap sympathetic neuronal cell bodies (Fig 2a-2c). This morphology can be observed in the Superior Cervical Ganglion (SCG) by postnatal day 2 (P2) (Fig 2a), a period of active sympathetic innervation of peripheral targets [36]. Over the course of the first eight postnatal weeks, sympathetic neurons increase in size (Fig 2b-d) as the neurons make target contacts and are exposed to target derived signals that promote cellular hypertrophy [37]. As neurons increase in size the number of neurons per unit area decreases, while the density of satellite glial cells remains constant over the first three postnatal weeks, with a decrease in glial density by 8 weeks (Fig 2e). This results in an increase in the number of glial cells associated with an individual neuron over the postnatal period.

**Fig 2.**
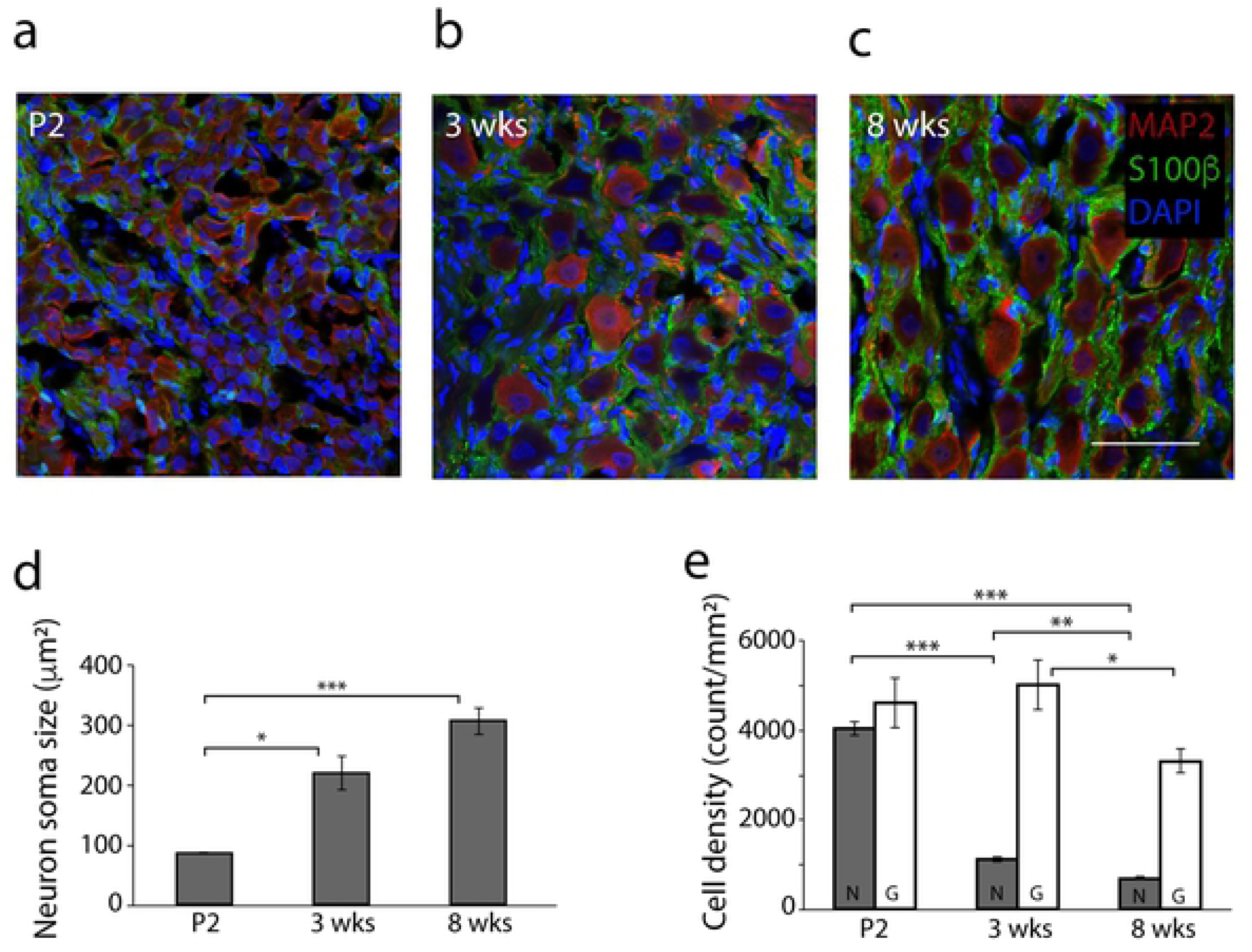
Postnatal development of neurons and satellite glia in the Superior Cervical Ganglion (SCG). (a-c) Representative confocal images of (a) P2 SCG, (b) 3 week (3 wks) old and (c) 8 weeks (8 wks) old SCG. Sympathetic neurons were stained for the neuron-specific marker MAP2 (red), satellite glial cells for the glial cell marker S100β (green) and cell nuclei using DAPI (blue). Scale bar = 60 µm. (d-e) Quantification of (d) neuron soma size, measured as average cell area in the section and (e) neuronal and glial cell densities from sections of P2 (n=3; mean ± s.e.m.), 3 wks (n=3; mean ± s.e.m.) and 8 wks animals (n=6; mean ± s.e.m.). ***p<0.001, **p<0.01, *p<0.05 determined by ANOVA followed by pairwise post hoc (Tukey’s HSD) comparison test.

### Satellite glia support survival and hypertrophy of cultured sympathetic neurons

We asked if satellite glial cells contributed to sympathetic neuron development in neonatal cultures by determining whether co-cultured glia acted to support the survival of NGF-deprived sympathetic neurons. Cultured sympathetic neurons are normally supported by the addition of 5 ng/ml NGF to the growth medium (Fig 3a-b). We did not observe a difference in neuron number in glial co-cultures compared to neurons grown alone. In contrast, NGF deprivation lead to almost complete neuronal cell death (Fig 3c, g) of neurons grown alone, while co-culture with satellite glia resulted in a partial rescue of neuronal survival (Fig 3d, g). We next asked if the survival effects of co-cultured glia were due to glia-derived NGF. NGF-deprived co-cultures of sympathetic neurons and satellite glia were treated with either an anti-NGF antibody to block endogenous NGF in the cultures or with K252a, a kinase inhibitor that blocks the TrkA receptor (Fig 3e, f and g). The survival effect of glial co-culture was abrogated following either treatment (Fig 3g), indicating that glial-produced neurotrophic factors can contribute to sympathetic neuron survival during the postnatal period.

**Fig 3.**
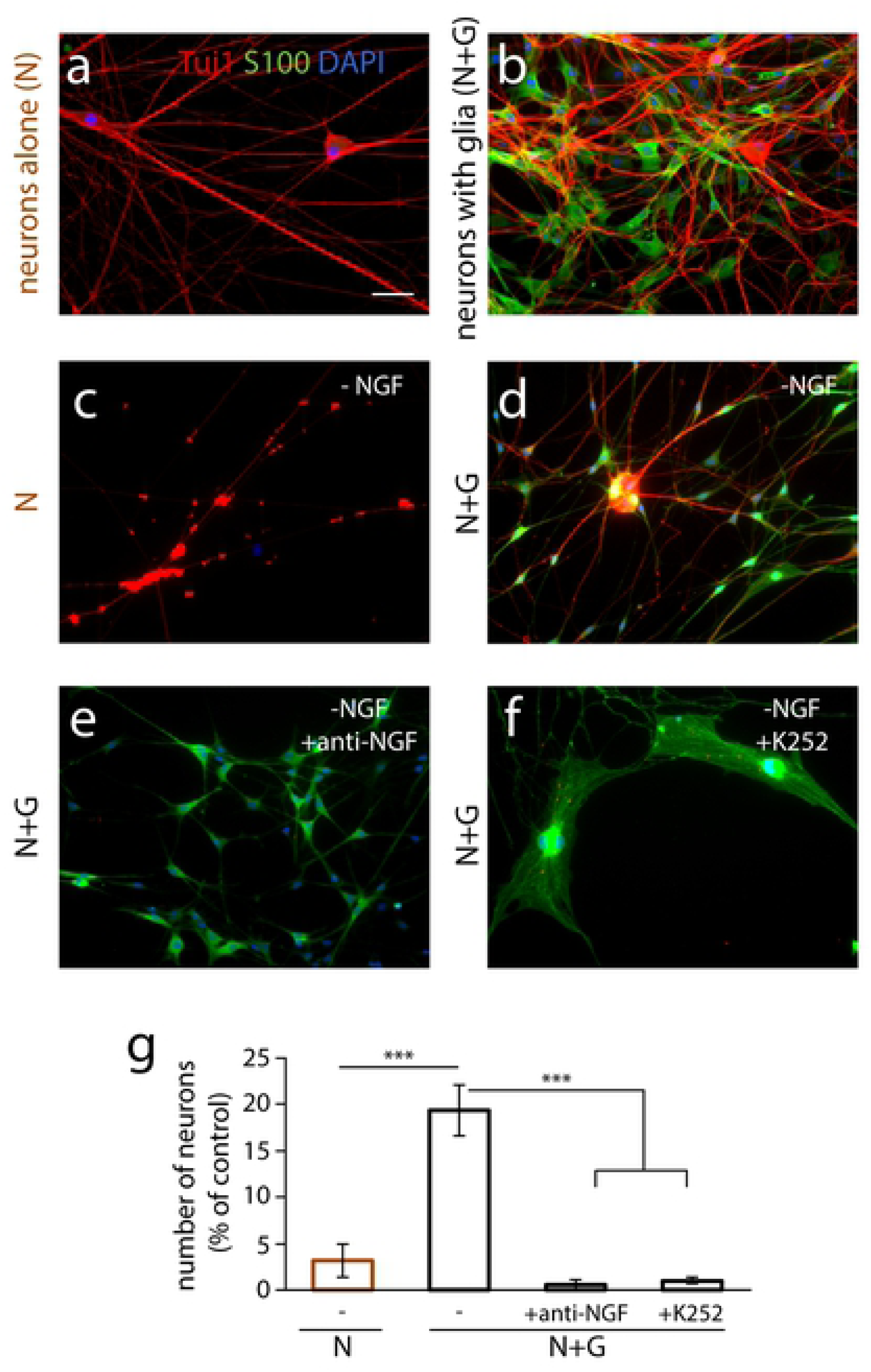
Satellite glial cells support neuronal survival. Satellite glial cells partly prevent sympathetic neuronal death upon NGF deprivation. (a-b) Establishment of sympathetic neuron-satellite glia co-cultures. Neurons (N) were cultured alone (a) or in the presence of satellite glia (b) in the presence of 5 ng/ml NGF in serum-containing medium. For NGF deprivation experiments (c-g) cultures were initially plated in the presence of 5 ng/ml NGF in serum-free medium and the medium was replaced after two days with NGF-free, serum-free medium. Cultures were fixed at 12 days *in vitro* (div) and stained for Tuj-1 (neuronal marker, in red), S100β (glial cell marker, in green) and DAPI (nuclear staining, in blue). Scale bar represents 50 µm. (c) Neurons alone (N), (d) Neurons and glia (N+G), (e) N+G with anti-NGF antibody (1:1000, final concentration 1 µg/ml), and (f) N+G with K252 (1:20000, final concentration 100 nM). (g) Quantification of cell survival upon NGF deprivation. Data are shown as percent neuronal survival compared to comparable cultures (neurons alone or neurons + glia) grown in the presence of 5 ng/ml NGF in serum-free medium (n = 3 independent cell culture experiments, One-way ANOVA, ***p<0.001). All data are represented as mean ± s.e.m.

We next asked if NGF protein could be detected by Western analysis of cell lysates prepared from isolated, cultured satellite glia. We found expression of NGF precursor forms (Fig 4a), supporting the idea that glia-derived NGF could have local effects on sympathetic neurons within the ganglion. We also detected expression of brain-derived neurotrophic factor, another member of the neurotrophin family, in cultured satellite glia (Fig 4b). Finally, we observed an increase in soma size for neurons grown in the presence of satellite glia (Table 1), suggesting a role for glia in neuronal hypertrophy during development. As expected, the cell area calculated for neurons in culture was larger than that seen for neuronal soma area in ganglionic sections (see Fig 2d), consistent with the constrained microenvironment *in vivo*.

**Fig 4.**
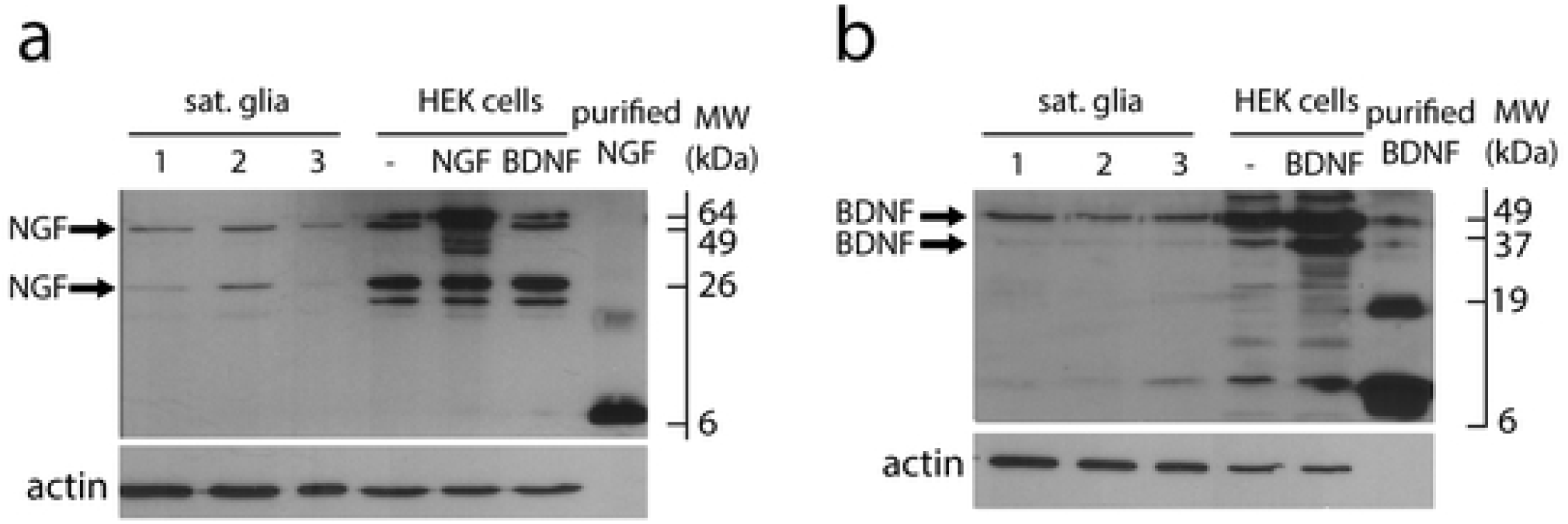
Satellite glial cells express neurotrophins. Western blot analyses reveal the expression of precursor forms of NGF (a) and BDNF (b) by satellite glial cells. Glia cell lysates were obtained from 3 independent cultures numbered 1-3. The specificity of the antibodies was assessed by comparing expression levels in HEK cells transfected with either empty (-), NGF-expressing (NGF) or BDNF-expressing (BDNF) plasmids. The mature purified neurotrophins were loaded as positive controls. β-actin expression was used as a loading control.

**Table 1.**
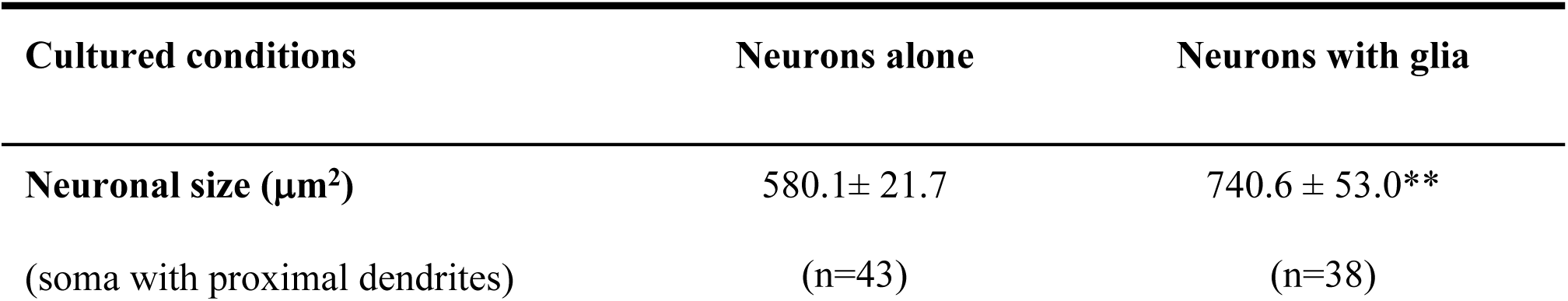
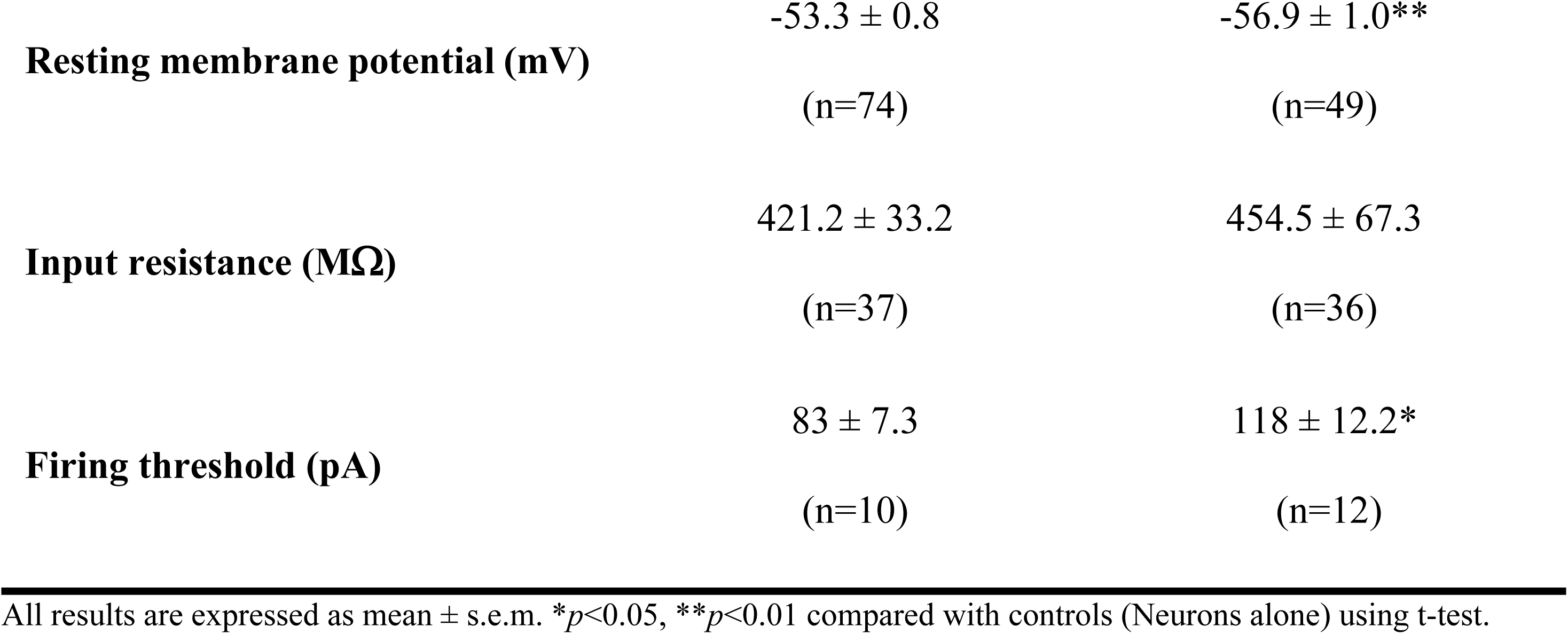
Neuronal characteristics of neurons co-cultured with or without glia.

### Satellite glia enhance spontaneous activity of cultured sympathetic neurons

We examined whether satellite glia influenced the active properties of sympathetic neurons and whether the effects were at the level of intrinsic neuronal firing properties and/or synaptic activity. Sympathetic neurons form nicotinic cholinergic synapses onto each other when in culture for 2 weeks or longer [38], providing a valuable and often used [36, 39, 40] cell culture model to study ganglionic cholinergic transmission between spinal preganglionic neurons and the postganglionic sympathetic neurons. We confirmed the cholinergic nature of sympathetic transmission in the presence of satellite glia by recording neuronal activity before and after infusion of the nicotinic cholinergic antagonist hexamethonium bromide. Total activity was measured for 20 minutes and quantified under control (0-3 min), hexamethonium (6-9 min) and washout (16-19 min) conditions. Activity was reduced to about 10% of control after hexamethonium infusion, showing a partial recovery to 55 % after 10 minutes of washout, indicative of the cholinergic nature of sympathetic transmission for neurons in the presence of satellite glia (Fig 5a-b).

**Fig 5.**
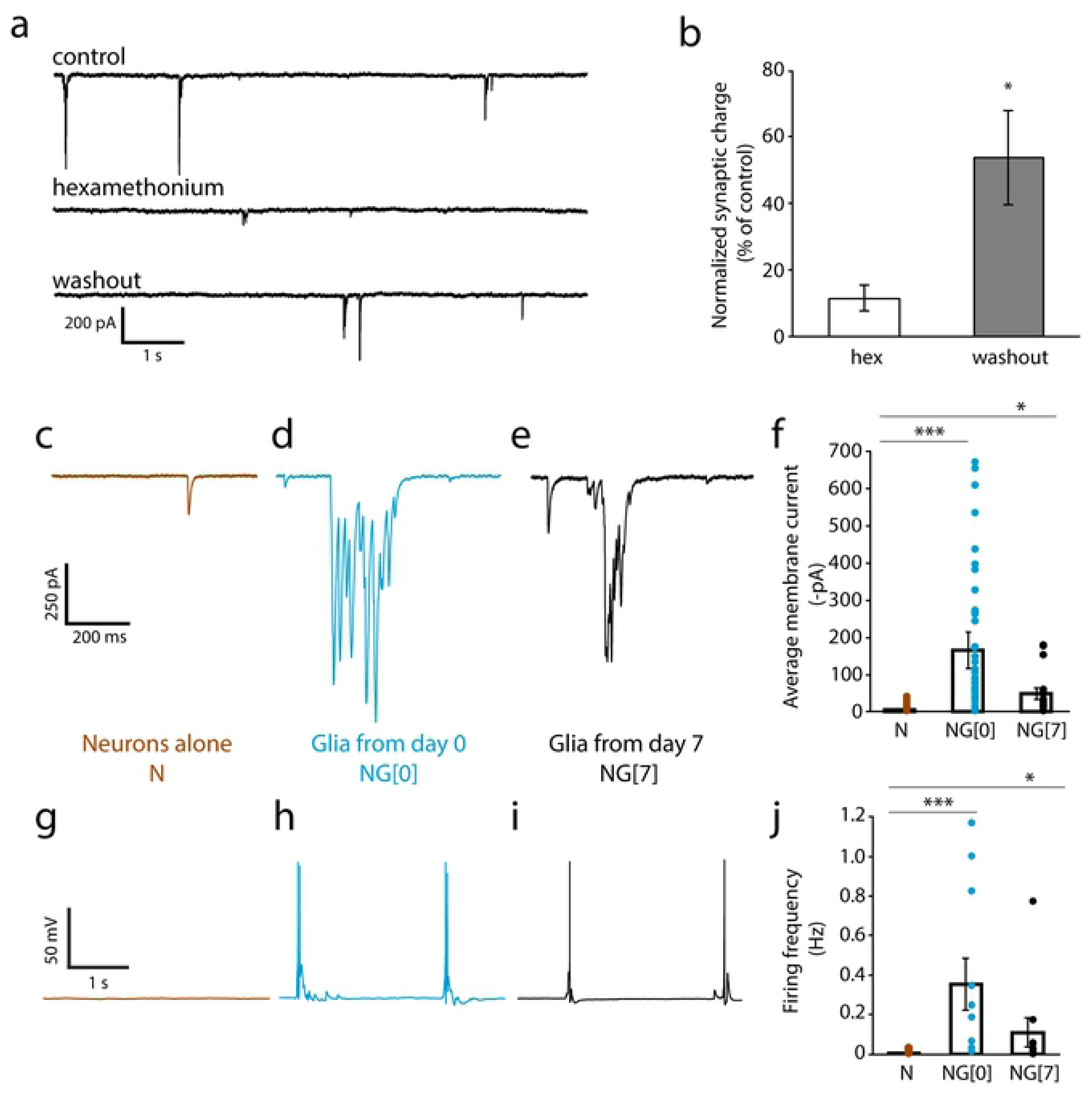
Satellite glial cells increase spontaneous activity of cultured sympathetic neurons. (a) Representative traces of spontaneous activity of neurons held in voltage-clamp at −60 mV, without hexamethonium, with 100 µM hexamethonium and after washout. (b) Average synaptic charge, normalized to control, for neurons treated with 100 µM hexamethonium (hex) and following washout (n=10 cells per condition; paired t-test; *p<0.05). (c-j) Neurons were cultured for 14 days alone (c, g) or in the presence of satellite glial cells starting from day 0 (d, h) or day 7 (e, i). NGF (5ng/ml) was included in all culture conditions to promote neuronal survival. (c-e) Representative voltage clamp traces showing that co-culture with satellite glial cells for the 14 days culture period, or for the last 7 days of the period increases current flow. (f) Quantification of synaptic activity. Total synaptic charge, defined as the area above the curve for neurons grown in the absence (Neurons alone, N) or presence of satellite glial cells for 14 days (NG[0]) or the last 7 days of the culture period (NG[7]), was quantified and average synaptic charge per 10 s duration was calculated. Plotted average membrane current values were quantified as averaged synaptic charge normalized to 1 ms duration. Therefore, the value of the average membrane current of e.g. −400 pA is equivalent to an average synaptic charge of 4 nC (n≥ 15 cells, Mann-Whitney U test, ***p<0.001, *p<0.05) (g-i) Representative current clamp traces showing that glial cells increase neuronal firing in cultured sympathetic neurons. (j) Quantification of neuronal firing rate in the absence (N) or presence of satellite glial cells for 14 (NG[0]) or 7 (NG[7]) days. (n≥10 cells, Mann-Whitney U test ***p<0.001, *p<0.05). Bars represent mean ± s.e.m.; dots represent data for individual cells.

We used this system to investigate the contribution of satellite glial cells to the development of sympathetic activity, measuring the spontaneous activity of sympathetic neurons cultured alone or in the presence of satellite glia. In the co-culture condition, neurons were grown with satellite glia for the full culture period (Glia from Day 0, NG[0]) as described in Methods. Spontaneous neuronal activity was recorded for 5 minutes for neurons in co-culture and neurons grown alone. We first assessed total current by recording in voltage clamp at a holding potential of - 60 mV. We found that the presence of glia resulted in a strong increase (>30 fold) in the total charge of sympathetic neurons when compared to the neuron alone condition (Fig 5c, d and f). Bursts of activity were commonly observed in the presence of glia, but were absent in the neuron alone cultures.

Generation of the post-mitotic neuron-alone cultures requires the use of AraC to block glia proliferation. We asked if the use of AraC in these cultures contributed to the low level of activity in comparison to the neuron-glia co-cultures, which were grown in the absence of AraC. We recorded neuronal activity in cultures (named “glia from day 7”, NG[7]) in which AraC was initially added to prevent glial proliferation; at day 7, satellite glial cells grown in a separate dish were re-plated on top of the neurons at approximately 100,000 cells per dish. These glia also formed a confluent layer by 10-14 div. The presence of glia from day 7 also increased total activity by about 10 fold (Fig 5e-f), indicating that glia still exert their effect on spontaneous activity when added at a later time point in culture even when the neurons had been exposed to AraC treatment.

We next asked whether the increase in neuronal activity was accompanied by an increase in the frequency of action potential firing. We recorded from neurons in current clamp and measured spontaneous neuronal activity, finding that neurons cultured in the presence of glial cells fired more action potentials than neurons cultured alone (Figs 5g-j). This increase in firing may underlie the occurrence of synchronous neurotransmitter release from pre-synaptic terminals and hence the bursts of activity seen in Figs 5d, 5e and 5f.

Overall, these results demonstrate that satellite glia derived from sympathetic ganglia increase the magnitude of neuronal inputs and the firing of cultured sympathetic neurons. Moreover, glia also exert their effects when added to the neuronal culture at a later stage (“glia from day 7”), suggesting that the effect is not dependent on early neurite extension and that glia act directly at synapses or at voltage-gated ion channels to enhance sympathetic activity.

### Excitable membrane properties of sympathetic neurons grown with satellite glia

We next investigated whether co-cultured glia affect neuronal firing properties in addition to synaptic properties. We examined the intrinsic membrane properties of neurons grown for 2-3 weeks alone or in the presence of satellite glia isolated from the same ganglia. We recorded from sympathetic neurons in whole cell current clamp and assessed resting membrane potential and input resistance (Table 1). There was no difference in neuronal input resistance between neurons grown in the presence or absence of glia. We observed a significant increase in neuronal resting membrane potential in the presence of glial cells.

We measured the effect of satellite glia on sympathetic intrinsic excitability by determining the neuronal firing in response to stimuli for neurons grown alone compared to neurons grown with glia. We analyzed the response of neurons to steps of depolarizing current in the presence of the cholinergic transmission blocker hexamethonium. Firing threshold, i.e. the minimum current needed to elicit an action potential, was determined by applying depolarizing currents steps in 10 pA increments (Fig 6a). We found a trend toward an increase in firing threshold in the presence of glia, although the difference was not statistically significant (Fig 6b). Next, we measured the number of spikes evoked by depolarizing current steps ranging from 0 to 400 pA (Fig 6c). There was a trend, that did not reach significance, towards a decrease in the number of APs fired by neurons cultured in the presence of glia in response to the current pulses (Fig 6d).

**Fig 6.**
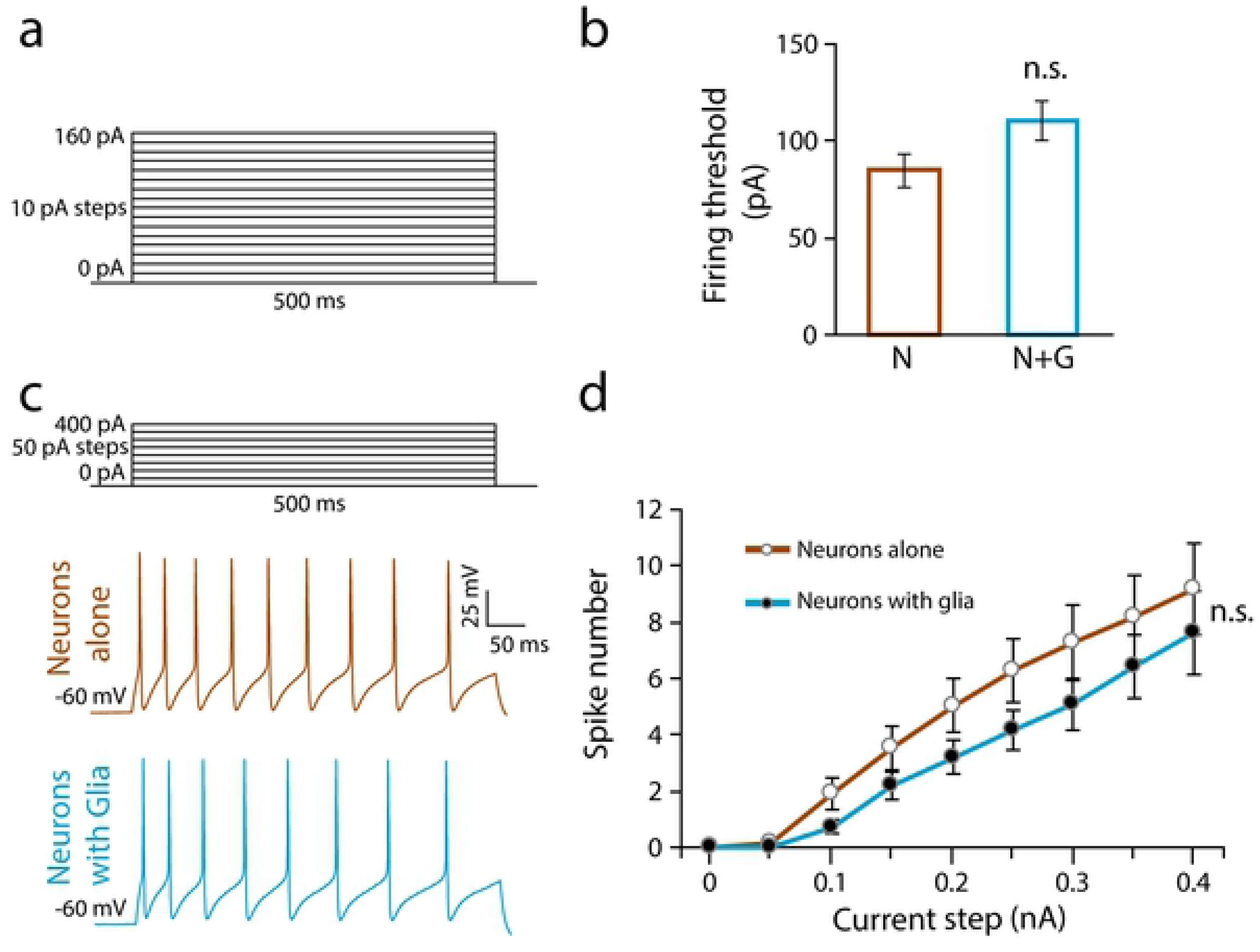
Satellite glial cells do not alter neuronal intrinsic excitability. (a) Illustration of the stimulus pattern applied to determine neuronal firing threshold. (b) Quantification of neuronal firing threshold between neurons alone (N) and neurons co-cultured with glia for 14 days (N+G) conditions. (Unpaired t-test, n ≥ 16 cells, n.s. not statistically different). (c) Illustration of the stimulus pattern applied to evoke action potential firing and representative neuronal traces in response to 400 pA current pulse for neurons grown for 14 days alone (brown, upper trace) or in the presence of glia (blue, lower trace). (d) Average number of action potentials evoked by current steps of increasing amplitude. (n≥16 cells, Mann-Whitney U test pairwise comparison for 400 pA current step, n.s. not statistically different). Results are represented as mean ± s.e.m.

### Satellite glia exert their effect via released factors

The effect of glia on sympathetic synaptic activity may be mediated by contact or by diffusible factors, or both. If diffusible factors play a role in this regulation, we would expect that glial cell-conditioned medium (GCM) would be sufficient to increase sympathetic neuron activity. We cultured glial cells until they reached confluency and allowed them to grow for an additional 3 days in serum-free medium before collecting the medium. The GCM was concentrated (see Methods) and added to sympathetic neurons that had been cultured alone for 7 days. Control medium (C) from fresh serum-free medium (not conditioned by glial cells) was concentrated to the same level as the GCM and added to neurons following the same protocol as for GCM (Figs. 7a and 7b). Following addition of GCM or C neurons were cultured for an additional 7 days. We compared spontaneous activity of neurons cultured in the presence of C or GCM at 14 div. The GCM did not affect neuronal survival, as neuron number was unaltered when compared to C (Fig 7c). Culture in GCM resulted in a >13 fold increase in sympathetic activity (Figs 7d-7f), comparable to the effect observed in the presence of satellite glial cells in culture from day 7 (Fig 5e-f). These data indicate that factor(s) released by satellite glial cells increase sympathetic activity. While we cannot rule out additional effects of glial cell contact, it seems likely that released factors are the main modulators of sympathetic activity, at least in cell culture, since GCM fully mimics the effect of satellite glial cells.

**Fig 7.**
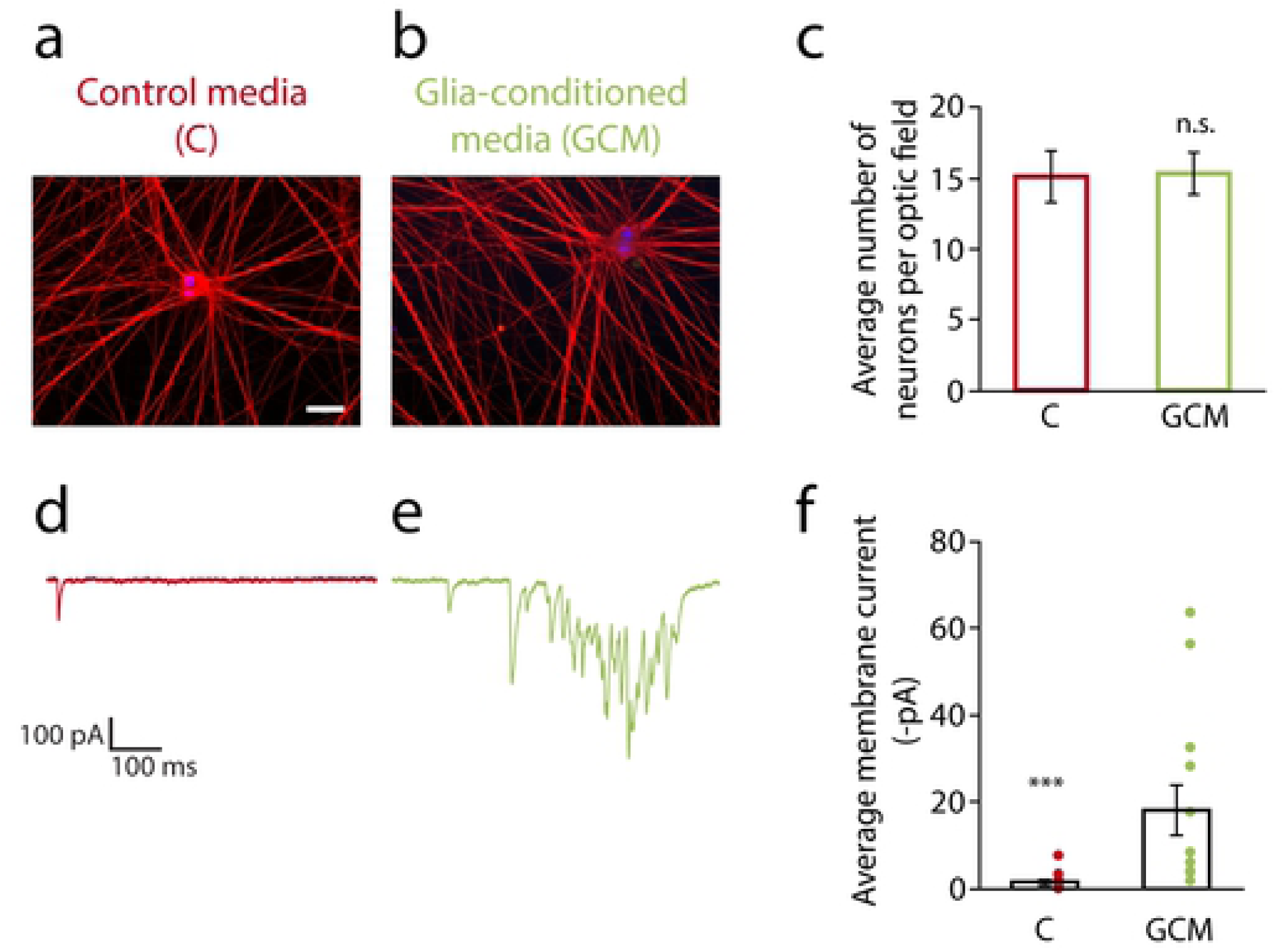
Glial cell-conditioned medium recapitulates the effect of satellite glial cells on cultured sympathetic neuron activity. (a-b) Neurons grown in control medium - C (a) or glial cell-conditioned medium - GCM (b) for the last 7 days of the 14-day culture period. NGF (5ng/ml) was added to both culture conditions to promote neuronal survival. Cells were fixed and stained for Tuj-1 (neuronal marker, in red), S100β (glial cell marker, in green) and DAPI (nuclear staining, in blue). Scale bar represents 50 µm. (c) Neuronal cell number was not altered by the cell culture condition. (n = 6 independent cell culture experiments, unpaired t-test, n.s.). (d-e) Representative voltage-clamp traces of neurons cultured in C (d) or GCM (e). (f) Quantification of average membrane current showing increased spontaneous activity in GCM (n ≥ 12 cells, non-parametric Mann-Whitney U test, ***p<0.001). Results are represented as mean ± s.e.m., dots represent data for individual cells.

We next asked whether glial cells had an effect on the development of synaptic sites. Soma and dendrites are important sites of synapse formation on peripheral sympathetic neurons [36, 41, 42]. We immunostained neurons using the vesicular acetylcholine transporter protein (VAChT) as a pre-synaptic marker, and the scaffold protein Shank (Shank) as a post-synaptic marker [43], and looked for their co-localization on cell bodies and proximal dendrites (Fig 8a). We quantified the number of pre-, post- and co-localized puncta and the size of co-localized puncta in cultures of neurons grown alone and in the presence of glial cells (Fig 8b-d). Co-culture with glia increased the number (Fig 8c) as well as the size (Fig 8d) of colocalized puncta on sympathetic neurons. There was a significant difference in the number of VAChT-positive puncta, but we observed no significant increase in the number of Shank-positive puncta (Fig 8b). This suggests a presynaptic effect of glia on the development of sympathetic synaptic sites, although additional postsynaptic mechanisms cannot be ruled out. These results suggest that glia promote increased sympathetic activity by promoting structural synapse formation.

**Fig 8.**
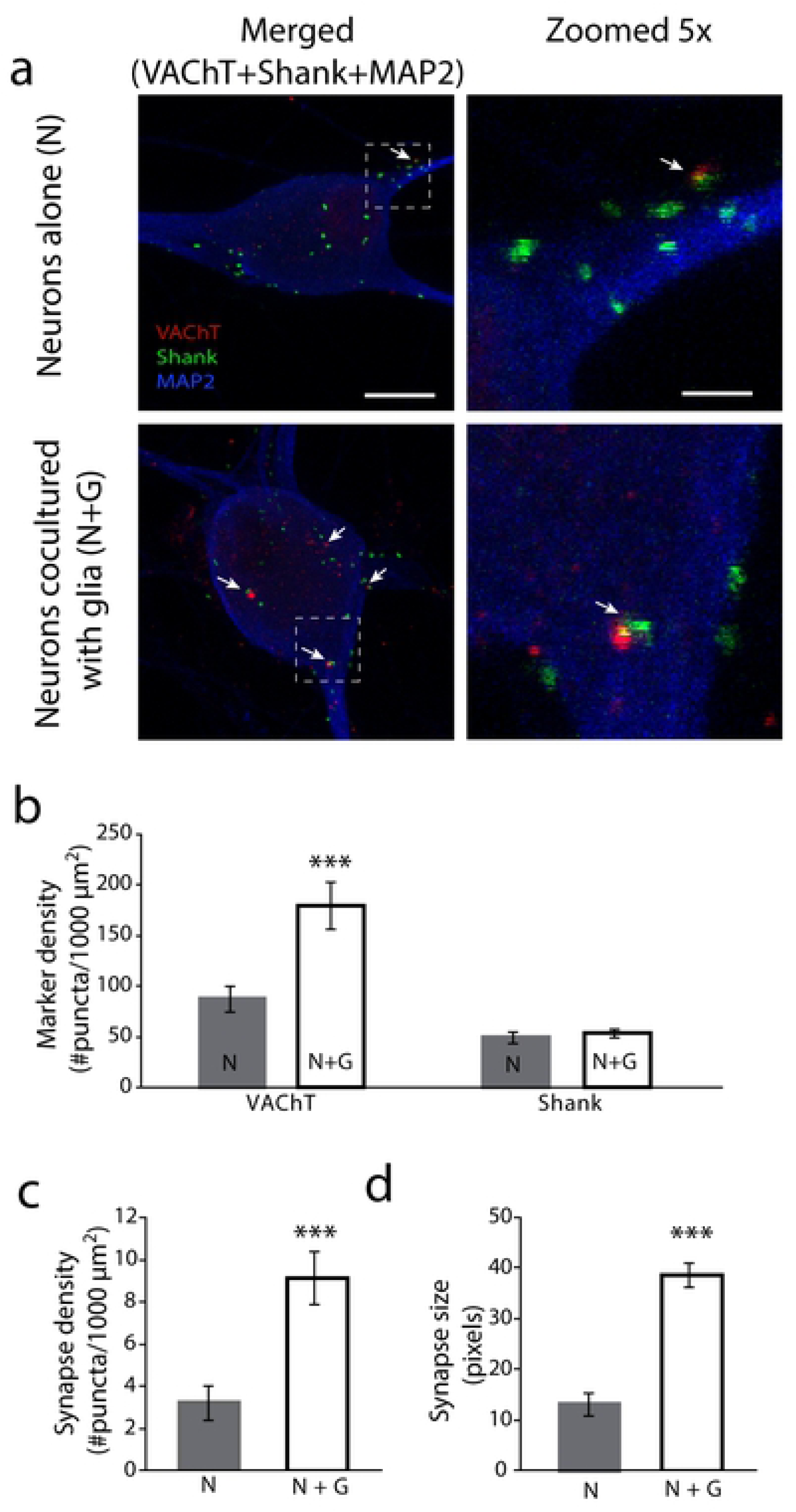
Satellite glial cells enhance cholinergic synapse formation. Neurons cultured in the absence (Neurons alone, N) or presence of satellite glial cells (N+G) were fixed, stained for synaptic markers, and analyzed by confocal microscopy. (a) Representative images of cells stained for the pre-synaptic marker VAChT (red), the post-synaptic marker Shank (green), and the dendritic marker MAP2 (blue). Boxed regions in left panels are magnified in the right panels to show colocalized puncta (arrows). Scale bar represents 10 µm in the left panels; 2 µm in the right panels. (b) Quantification of VAChT and Shank puncta on sympathetic neuronal soma and proximal dendrites showing a glia-dependent increase in the expression of VAChT, but not Shank-containing puncta (n ≥ 61 cells, unpaired t-test ***p<0.001 and n.s., respectively). Quantification of co-localized synaptic puncta density (c) and size (d) on neuronal soma and proximal dendrites showing that glia induce structural synapse formation (n ≥ 61 cells, unpaired t-test, ***p<0.001, **p<0.01, respectively).

## Discussion

We report a role for satellite glial cells in the establishment of mature sympathetic neuron structure and function within peripheral ganglia. Satellite glia contribute to the survival of cultured postnatal sympathetic neurons and potentiate sympathetic cholinergic synaptic activity and structural synapse formation. The effects of satellite glia are mediated by released factors, which include NGF and BDNF, two known modulators of sympathetic neuron activity [39, 44, 45]. This work defines sympathetic satellite glia as regulators of peripheral neuronal development and provides a new path for understanding mechanisms leading to heightened sympathetic tone.

The actions of satellite glial cells in regulating synapse formation and neuronal activity within the sympathetic ganglia share some common features with astrocytes in the central nervous system [1, 46]. While this illustrates a convergence in function between these two glial cell types of different embryonic origins, the neuronal targets of glial regulation are distinct. Astrocytes regulate many aspects of excitatory glutamatergic and inhibitory GABAergic transmission in the CNS [13, 47–49], while in the periphery, we show that sympathetic satellite glia promote the development of spontaneous network activity at cholinergic synapses.

Outside of the mammalian CNS, glial regulation of cholinergic systems has been reported in *Lymnaea stagnalis*, where cholinergic neurons grown in the presence of glial cells have decreased postsynaptic responses to presynaptic stimulation [50]. Non-neuronal ganglionic cells also regulate short-term plasticity at sympathetic cholinergic autapses without an effect on synaptic development [29]. Earlier work suggested a role in synapse formation by showing that unidentified non-neuronal cells promoted evoked release of acetylcholine in mass cultures of sympathetic neurons [51]. Taken together with our findings of increased synapse number and spontaneous synaptic transmission in sympathetic neurons co-cultured with satellite glial cells, these studies show that glial modulation of cholinergic properties is characterized by system-specific properties. Our work shows that within the developing sympathetic system, glial cells release soluble factors that contribute to the development and dynamics of cholinergic circuits.

System-specific characteristics of glial modulation are also seen by comparing satellite glial actions in peripheral sensory and sympathetic ganglia [23]. Satellite glial cells of the sensory ganglia have been studied in the context of abnormal pain conditions and were found to contribute to neuronal hyperexcitability [52, 53]. In our study we analyzed, and did not find a significant difference in intrinsic excitability of sympathetic neurons cultured in the presence of satellite glia (Fig 5). This differential effect of satellite glia in sympathetic and sensory ganglia may reflect anatomical differences between these ganglia, since sensory ganglia do not receive inputs from central preganglionic neurons, and do not contain dendrites or synapses. It thus seems likely that glia affect distinct neuronal properties in these two peripheral ganglia, an idea that is supported by the synaptic effects and absence of changes in excitability in our cultures.

Our finding of glial regulation of cholinergic transmission and presynaptic protein expression suggests a regulatory circuit in which glial factors act to increase neuronal acetylcholine release, which in turn acts on the glial cells to modulate glial activity. This model is supported by recent work demonstrating changes in sympathetic satellite glial activity in response to glial muscarinic cholinergic receptor activation [54]. This cholinergic signaling resulted in an increase in glial calcium signaling, glial activation and electrical coupling between glial cells, suggesting that activity in neural circuits may be set by reciprocal signaling between neurons and their surrounding glia.

In the CNS, astrocyte cell function is also modulated by cholinergic signaling, resulting in glial regulation of glutamatergic or GABAergic neurotransmission. In the hippocampus, for instance, astrocytes act as a sensor for septal-derived acetylcholine associated with wakefulness, resulting in gating of glutamatergic transmission through NMDA receptors [46]. Cholinergic modulation of hippocampal astrocytes also leads to long-term inhibition of dentate granule cells through direct glial excitation of inhibitory interneurons [55]. In addition, cholinergic modulation of glial cell function has been observed in the retina [56] and the enteric nervous system [57]. Together, these studies demonstrate wide-spread actions of cholinergic signaling on glial activity states; however less is known about reciprocal signaling from astrocytes to cholinergic synapses. Our work in the peripheral nervous system suggests that these effects may be part of a broader regulatory system that includes glial control of their cholinergic inputs.

Satellite glia regulated multiple developmental processes in this study, including neuronal survival (Fig 3), cell body hypertrophy (Table 1) and cholinergic synapse formation (Fig 7). These are all promoted by NGF in developing sympathetic neurons [36, 58–60]. We showed that NGF released by satellite glia partially supported the survival of the cultured sympathetic neurons. The production of neurotrophins by satellite glia is consistent with the reported expression of neurotrophins in other central and peripheral glial populations [61–65]. However, extensive evidence points to the central role of target-derived NGF produced by peripheral organs in the survival and morphological maturation of postnatal sympathetic neurons [37, 66]. Thus, our data suggest that glial-derived neurotrophic factors may provide a secondary source of neurotrophic signaling during postnatal development and in the mature the sympathetic circuit.

We previously showed synaptic modulation of sympathetic cholinergic transmission by neurotrophins [39], but further work will be needed to determine if glial-derived neurotrophins contribute to the synaptic effects of glial cells. We do not expect that glial-derived NGF is the primary source of neurotrophic signaling in this system, as NGF is retrogradely transported from peripheral targets *in vivo* [67]. It is interesting to speculate, however, that ganglionic sources of such factors could play a stabilizing role during development or following nerve injury. Peripheral nerve injury is accompanied by a reduction in NGF retrograde transport [68] and a dramatic decrease in sympathetic neuron activity and cholinergic synapses within the ganglia [69]. Thus, ganglionic sources of NGF could provide a compensatory source of neurotrophic signaling and would be consistent with activation of glia during pathological disruptions [70, 71].

The effects of satellite glia on sympathetic synaptic function suggest the potential for ongoing glial regulation in the sympathetic system. This is of particular interest in pathological situations, such as in cardiovascular disorders in which sympathetic over-activation is a common feature [19–21]. This idea is consistent with recently published work using selective activation of a glial-expressed Gq protein-coupled receptor in transgenic mice to show that acutely activated glial cells *in vivo* increased heart rate and cardiac output through the actions of the peripheral sympathetic system [27, 28]. This increase in heart rate was abolished by selective inhibition of peripheral glia activation, further establishing satellite glia as regulators of sympathetic-mediated cardiac function. The work described here demonstrates a link between ganglionic satellite glia and functional changes in the electrical properties of sympathetic neurons, providing a mechanistic model for the actions of satellite glia in driving heightened sympathetic tone and suggesting these glia as potential new targets to treat diseases of the peripheral organs.

## Acknowledgements

We thank the members of the Turrigiano laboratory for their support and discussions on the project, and S. Van Hooser and J. Maier for assistance with coding in MatLab. We gratefully acknowledge G. Turrigiano, V. Tatarvarty, and M. Nahmani for help in refining the manuscript. We also thank Cagla Eroglu for providing us the Puncta Analyzer plug-in, and Masami Kojima for providing plasmids.

## References

1. Allen NJ, Barres BA. Neuroscience: Glia - more than just brain glue. Nature. 2009;457(7230):675–7.

2. Freeman MR, Rowitch DH. Evolving concepts of gliogenesis: a look way back and ahead to the next 25 years. Neuron. 2013;80(3):613–23.

3. Devinsky O, Vezzani A, Najjar S, De Lanerolle NC, Rogawski MA. Glia and epilepsy: excitability and inflammation. Trends Neurosci. 2013;36(3):174–84.

4. Araque A, Carmignoto G, Haydon PG, Oliet SH, Robitaille R, Volterra A. Gliotransmitters travel in time and space. Neuron. 2014;81(4):728–39.

5. Pannasch U, Rouach N. Emerging role for astroglial networks in information processing: from synapse to behavior. Trends Neurosci. 2013;36(7):405–17.

6. Ullian EM, Sapperstein SK, Christopherson KS, Barres BA. Control of synapse number by glia. Science. 2001;291(5504):657–61.

7. Buard I, Steinmetz CC, Claudepierre T, Pfrieger FW. Glial cells promote dendrite formation and the reception of synaptic input in Purkinje cells from postnatal mice. Glia. 2010;58(5):538–45.

8. Hughes EG, Elmariah SB, Balice-Gordon RJ. Astrocyte secreted proteins selectively increase hippocampal GABAergic axon length, branching, and synaptogenesis. Mol Cell Neurosci. 2010;43(1):136–45.

9. Allen NJ, Bennett ML, Foo LC, Wang GX, Chakraborty C, Smith SJ, et al. Astrocyte glypicans 4 and 6 promote formation of excitatory synapses via GluA1 AMPA receptors. Nature. 2012;486(7403):410–4.

10. Pfrieger FW, Barres BA. Synaptic efficacy enhanced by glial cells in vitro. Science. 1997;277(5332):1684–7.

11. Pannasch U, Freche D, Dallerac G, Ghezali G, Escartin C, Ezan P, et al. Connexin 30 sets synaptic strength by controlling astroglial synapse invasion. Nat Neurosci. 2014;17(4):549–58.

12. Chung WS, Clarke LE, Wang GX, Stafford BK, Sher A, Chakraborty C, et al. Astrocytes mediate synapse elimination through MEGF10 and MERTK pathways. Nature. 2013;504(7480):394–400.

13. Clarke LE, Barres BA. Emerging roles of astrocytes in neural circuit development. Nat Rev Neurosci. 2013;14(5):311–21.

14. Allen NJ. Role of glia in developmental synapse formation. Current opinion in neurobiology. 2013;23(6):1027–33.

15. Corty MM, Freeman MR. Cell biology in neuroscience: Architects in neural circuit design: glia control neuron numbers and connectivity. The Journal of cell biology. 2013;203(3):395.

16. Garden GA, La Spada AR. Intercellular (mis)communication in neurodegenerative disease. Neuron. 2012;73(5):886–901.

17. McGann JC, Lioy DT, Mandel G. Astrocytes conspire with neurons during progression of neurological disease. Current opinion in neurobiology. 2012;22(5):850–8.

18. Barman SM, Yates BJ. Deciphering the Neural Control of Sympathetic Nerve Activity: Status Report and Directions for Future Research. Frontiers in Neuroscience. 2017;11(730):1–14.

19. Guyenet PG. The sympathetic control of blood pressure. Nat Rev Neurosci. 2006;7(5):335–46.

20. Kaye D, Esler M. Sympathetic neuronal regulation of the heart in aging and heart failure. Cardiovasc Res. 2005;66(2):256–64.

21. Dutta P, Courties G, Wei Y, Leuschner F, Gorbatov R, Robbins CS, et al. Myocardial infarction accelerates atherosclerosis. Nature. 2012;487(7407):325–9.

22. Saper CB. The central autonomic nervous system: conscious visceral perception and autonomic pattern generation. Annual review of neuroscience. 2002;25:433–69.

23. Hanani M. Satellite glial cells in sympathetic and parasympathetic ganglia: in search of function. Brain Res Rev. 2010;64(2):304–27.

24. Bushong EA, Martone ME, Jones YZ, Ellisman MH. Protoplasmic astrocytes in CA1 stratum radiatum occupy separate anatomical domains. Journal of Neuroscience; 2002;22(1):183–92.

25. Huang TY, Hanani M, Ledda M, De Palo S, Pannese E. Aging is associated with an increase in dye coupling and in gap junction number in satellite glial cells of murine dorsal root ganglia. Neuroscience. 2006;137(4):1185–92.

26. Hu P, McLachlan EM. Inflammation in sympathetic ganglia proximal to sciatic nerve transection in rats. Neuroscience letters. 2004;365(1):39–42.

27. Agulhon C, Boyt KM, Xie AX, Friocourt F, Roth BL, McCarthy K. Modulation of the autonomic nervous system by acute glial cell Gq-GPCR activation in vivo. The Journal of physiology. 2013;591(22):5599–609.

28. Xie AX, Lee JJ, McCarthy KD. Ganglionic GFAP glial Gq-GPCR signaling enhances heart functions in vivo. JCI insight. 2017;2(2):e90565.

29. Perez-Gonzalez AP, Albrecht D, Blasi J, Llobet A. Schwann cells modulate short-term plasticity of cholinergic autaptic synapses. The Journal of physiology. 2008;586(Pt 19):4675–91.

30. Verdi JM, Groves AK, Farinas I, Jones K, Marchionni MA, Reichardt LF, et al. A reciprocal cell-cell interaction mediated by NT-3 and neuregulins controls the early survival and development of sympathetic neuroblasts. Neuron. 1996;16(3):515–27.

31. Tropea M, Johnson MI, Higgins D. Glial cells promote dendritic development in rat sympathetic neurons in vitro. Glia. 1988;1(6):380–92.

32. Barish ME. Modulation of the electrical differentiation of neurons by interactions with glia and other non-neuronal cells. Perspectives on developmental neurobiology. 1995;2(4):357–70.

33. McFarlane S, Cooper E. Extrinsic factors influence the expression of voltage-gated K currents on neonatal rat sympathetic neurons. Journal of Neuroscience; 1993;13(6):2591–600.

34. Suter U, Heymach JV, Jr., Shooter EM. Two conserved domains in the NGF propeptide are necessary and sufficient for the biosynthesis of correctly processed and biologically active NGF. The EMBO journal. 1991;10(9):2395–400.

35. Heymach JV, Jr., Shooter EM. The biosynthesis of neurotrophin heterodimers by transfected mammalian cells. The Journal of biological chemistry. 1995;270(20):12297–304.

36. Sharma N, Deppmann CD, Harrington AW, St Hillaire C, Chen ZY, Lee FS, et al. Long-distance control of synapse assembly by target-derived NGF. Neuron. 2010;67(3):422–34.

37. Davis BM, Wang HS, Albers KM, Carlson SL, Goodness TP, McKinnon D. Effects of NGF overexpression on anatomical and physiological properties of sympathetic postganglionic neurons. Brain research. 1996;724(1):47–54.

38. O’Lague PH, Obata K, Claude P, Furshpan EJ, Potter DD. Evidence for cholinergic synapses between dissociated rat sympathetic neurons in cell culture. Proc Natl Acad Sci U S A. 1974;71(9):3602–6.

39. Luther JA, Enes J, Birren SJ. Neurotrophins regulate cholinergic synaptic transmission in cultured rat sympathetic neurons through a p75-dependent mechanism. J Neurophysiol. 2013;109(2):485–96.

40. Campanucci V, Krishnaswamy A, Cooper E. Diabetes depresses synaptic transmission in sympathetic ganglia by inactivating nAChRs through a conserved intracellular cysteine residue. Neuron. 2010;66(6):827–34.

41. Gingras J, Rassadi S, Cooper E, Ferns M. Agrin plays an organizing role in the formation of sympathetic synapses. J Cell Biol. 2002;158(6):1109–18.

42. Gingras J, Ferns M. Expression and localization of agrin during sympathetic synapse formation in vitro. J Neurobiol. 2001;48(3):228–42.

43. Parker MJ, Zhao S, Bredt DS, Sanes JR, Feng G. PSD93 regulates synaptic stability at neuronal cholinergic synapses. Journal of Neuroscience; 2004;24(2):378–88.

44. Slonimsky JD, Yang B, Hinterneder JM, Nokes EB, Birren SJ. BDNF and CNTF regulate cholinergic properties of sympathetic neurons through independent mechanisms. Molecular and Cellular Neuroscience. 2003;23(4):648–60.

45. Arias ER, Valle-Leija P, Morales MA, Cifuentes F. Differential contribution of BDNF and NGF to long-term potentiation in the superior cervical ganglion of the rat. Neuropharmacology. 2014;81:206–14.

46. Papouin T, Dunphy J, Tolman M, Foley JC, Haydon PG. Astrocytic control of synaptic function. Philosophical transactions of the Royal Society of London Series B, Biological sciences. 2017;372(1715).

47. Baldwin KT, Eroglu C. Molecular mechanisms of astrocyte-induced synaptogenesis. Current opinion in neurobiology. 2017;45:113–20.

48. Um JW. Roles of Glial Cells in Sculpting Inhibitory Synapses and Neural Circuits. Frontiers in molecular neuroscience. 2017;10:381.

49. Turko P, Groberman K, Browa F, Cobb S, Vida I. Differential Dependence of GABAergic and Glutamatergic Neurons on Glia for the Establishment of Synaptic Transmission. Cerebral cortex (New York, NY : 1991). 2019;29(3):1230–43.

50. Smit AB, Syed NI, Schaap D, van Minnen J, Klumperman J, Kits KS, et al. A glia-derived acetylcholine-binding protein that modulates synaptic transmission. Nature. 2001;411(6835):261–8.

51. O’Lague PH, Furshpan EJ, Potter DD. Studies on rat sympathetic neurons developing in cell culture. II. Synaptic mechanisms. Developmental biology. 1978;67(2):404–23.

52. Hanani M. Satellite glial cells in sensory ganglia: from form to function. Brain Res Brain Res Rev. 2005;48(3):457–76.

53. Retamal MA, Alcayaga J, Verdugo CA, Bultynck G, Leybaert L, Saez PJ, et al. Opening of pannexin- and connexin-based channels increases the excitability of nodose ganglion sensory neurons. Front Cell Neurosci. 2014;8(158):1–12.

54. Feldman-Goriachnik R, Wu B, Hanani M. Cholinergic responses of satellite glial cells in the superior cervical ganglia. Neuroscience letters. 2018;671:19–24.

55. Pabst M, Braganza O, Dannenberg H, Hu W, Pothmann L, Rosen J, et al. Astrocyte Intermediaries of Septal Cholinergic Modulation in the Hippocampus. Neuron. 2016;90(4):853–65.

56. Rosa JM, Bos R, Sack GS, Fortuny C, Agarwal A, Bergles DE, et al. Neuron-glia signaling in developing retina mediated by neurotransmitter spillover. eLife. 2015;4.

57. Delvalle NM, Fried DE, Rivera-Lopez G, Gaudette L, Gulbransen BD. Cholinergic activation of enteric glia is a physiological mechanism that contributes to the regulation of gastrointestinal motility. American journal of physiology Gastrointestinal and liver physiology. 2018;315(4):G473–g83.

58. Ruit KG, Osborne PA, Schmidt RE, Johnson EM, Jr., Snider WD. Nerve growth factor regulates sympathetic ganglion cell morphology and survival in the adult mouse. Journal of Neuroscience; 1990;10(7):2412–9.

59. Kuruvilla R, Zweifel LS, Glebova NO, Lonze BE, Valdez G, Ye H, et al. A neurotrophin signaling cascade coordinates sympathetic neuron development through differential control of TrkA trafficking and retrograde signaling. Cell. 2004;118(2):243–55.

60. Lockhart ST, Mead JN, Pisano JM, Slonimsky JD, Birren SJ. Nerve growth factor collaborates with myocyte-derived factors to promote development of presynaptic sites in cultured sympathetic neurons. J Neurobiol. 2000;42(4):460–76.

61. Rudge JS, Alderson RF, Pasnikowski E, McClain J, Ip NY, Lindsay RM. Expression of Ciliary Neurotrophic Factor and the Neurotrophins-Nerve Growth Factor, Brain-Derived Neurotrophic Factor and Neurotrophin 3-in Cultured Rat Hippocampal Astrocytes. The European journal of neuroscience. 1992;4(6):459–71.

62. Wetmore C, Olson L. Neuronal and nonneuronal expression of neurotrophins and their receptors in sensory and sympathetic ganglia suggest new intercellular trophic interactions. J Comp Neurol. 1995;353(1):143–59.

63. Jha MK, Kim JH, Song GJ, Lee WH, Lee IK, Lee HW, et al. Functional dissection of astrocyte-secreted proteins: Implications in brain health and diseases. Progress in neurobiology. 2018;162:37–69.

64. Thippeswamy T, McKay JS, Morris R, Quinn J, Wong LF, Murphy D. Glial-mediated neuroprotection: Evidence for the protective role of the NO-cGMP pathway via neuron-glial communication in the peripheral nervous system. Glia. 2005;49(2):197–210.

65. Elmariah SB, Hughes EG, Oh EJ, Balice-Gordon RJ. Neurotrophin signaling among neurons and glia during formation of tripartite synapses. Neuron Glia Biol. 2004;1(4):1–11.

66. Fagan AM, Zhang H, Landis S, Smeyne RJ, Silos-Santiago I, Barbacid M. TrkA, but not TrkC, receptors are essential for survival of sympathetic neurons in vivo. The Journal of Neuroscience; 1996;16(19):6208–18.

67. Harrington AW, Ginty DD. Long-distance retrograde neurotrophic factor signalling in neurons. Nat Rev Neurosci. 2013;14(3):177–87.

68. Raivich G, Hellweg R, Kreutzberg GW. NGF receptor-mediated reduction in axonal NGF uptake and retrograde transport following sciatic nerve injury and during regeneration. Neuron. 1991;7(1):151–64.

69. Purves D. Functional and structural changes in mammalian sympathetic neurones following interruption of their axons. The Journal of physiology. 1975;252(2):429–63.

70. Zhou XF, Deng YS, Chie E, Xue Q, Zhong JH, McLachlan EM, et al. Satellite-cell-derived nerve growth factor and neurotrophin-3 are involved in noradrenergic sprouting in the dorsal root ganglia following peripheral nerve injury in the rat. The European journal of neuroscience. 1999;11(5):1711–22.

71. Jones EV, Bouvier DS. Astrocyte-secreted matricellular proteins in CNS remodelling during development and disease. Neural Plast. 2014;2014(321209):1–12.

